# Assessing the molecular and phenotypic contribution of a horizontally acquired region to yeast adaptation

**DOI:** 10.1101/2025.11.11.687820

**Authors:** Andrés Romero, Camila Bastías, Matteo De Chiara, Xanita Saayman, Hajar Cherkaoui, Benjamin Barré, Francisco A. Cubillos, Claudio Martinez, Eduardo I. Kessi-Pérez, Gianni Liti, Francisco Salinas

## Abstract

Horizontal Gene Transfer (HGT) is the movement of genetic material across species. In *Saccharomyces cerevisiae*, a DNA segment known as Region B was acquired horizontally from a distant yeast species. This region (∼17 Kb) encodes 5 genes and is present in the genomes of yeast strains from different phylogenetic clades, with its contribution to yeast niche-specific adaptation remaining unclear. In this work, the genomic structure of Region B was analyzed in yeast strains from the ScRAP (*Saccharomyces cerevisiae* Reference Assembly Panel) collection, identifying 10 variants that maintain a circular continuity. To assess the role of Region B in yeast adaptation, we performed a high-throughput phenotyping of the ScRAP collection under different growth conditions, identifying that Region B is potentially associated with higher tolerance to oxidative stress. Then, we selected a single yeast strain from the ScRAP collection for characterization of the transcriptional activity of each gene within Region B using a fluorescent reporter. The results revealed that gene expression depends on the host’s genetic background and transcription factors encoded within Region B. To identify the genetic determinants involved in Region B expression within different genetic backgrounds, three expression Quantitative Trait Loci (eQTLs) were mapped and validated. Finally, by performing Region B deletion, we determine the contribution of this region to different fermentative phenotypes, including fermentation rate and amino acid consumption. Altogether, our results suggest a complex regulatory interaction between the horizontally acquired genes and the host genome that contributes to yeast adaptation under fermentation conditions.

## 1. Introduction

The budding yeast *Saccharomyces cerevisiae* is a model eukaryotic organism, and a biotechnology platform associated with human activities, dating back over 9,000 years to early intentional fermentations of rice, fruits, and honey (McGovern, et al. 2004). After the publication of the reference yeast genome (Goffeau, et al. 1996), a growing number of studies have explored the genetic and phenotypic diversity of *S. cerevisiae* strains isolated from different environments, including human-associated niches such as fermented beverages and food (Liti, et al. 2009; Gallone, et al. 2016; Gonçalves, et al. 2016; Legras, et al. 2018; Peter, et al. 2018; O’Donnell, et al. 2023). Notably, the release of 1,011 genomes derived from worldwide *S. cerevisiae* isolates provided a comprehensive view of the genetic diversity in the species (Peter, et al. 2018). This collection established that *S. cerevisiae* pangenome harbors approximately 7,800 non-redundant Open Reading Frames (ORFs), among which about 2,800 ORFs comprise the variable genome (Peter, et al. 2018). The quality of yeast genome assemblies was increased with long-read sequencing technologies, generating the *S. cerevisiae* Reference Assembly Panel (ScRAP) collection, which comprises 142 telomere-to-telomere genomes from yeast strains isolated from wild and human-related environments (O’Donnell, et al. 2023). Recently, high-quality assemblies for 1089 yeast genomes were obtained using a long-read technology, determining the genetic basis of molecular and organismal traits with unprecedented resolution (Loegler, et al. 2025).

In eukaryotes, Horizontal Gene Transfer (HGT) occurs infrequently due to cellular barriers that prevent the incorporation of foreign genetic material. In yeast, HGT constitutes approximately 2.3% of the variable genome (Peter, et al. 2018), with putative functions related to oxidative stress, anaerobic growth, pH tolerance, and carbon and nitrogen metabolism. Besides, multiple studies have demonstrated the contribution of horizontally acquired genes to enhance yeast fitness in fermentative environments (Novo, et al. 2009; Galeote, et al. 2010; Wenger, et al. 2010; Marsit, et al. 2015; Marsit, et al. 2016; Legras, et al. 2018; Devia, et al. 2020; Figueroa, et al. 2025). Initially, three horizontally acquired regions were described in the EC1118 wine yeast strain, corresponding to Region A (∼38 Kb), Region B (∼17Kb), and Region C (∼65 Kb) (Novo, et al. 2009). These regions were acquired from different species of the *Torulaspora* genus for Regions A and C, and *Zygosaccharomyces parabailii* for Region B; species commonly found cohabiting with *S. cerevisiae* during the early stages of grape fermentation (Novo, et al. 2009; Peter, et al. 2018; O’Donnell, et al. 2023). To date, only four genes belonging to Region C have been functionally characterized, showing important roles in wine fermentation, consistent with the almost exclusive presence of this region in isolates from the Wine-European clade (Galeote, et al. 2010; Wenger, et al. 2010; Marsit, et al. 2015; Marsit, et al. 2016; Legras, et al. 2018; Peter, et al. 2018; O’Donnell, et al. 2023). Recently, Region A was found to be related to wine fermentation, contributing to fermentation completion under nitrogen-limited conditions (Figueroa, et al. 2025). Altogether, beyond the mentioned examples, the role of Region B in yeast adaptation remains largely uncharted this far.

Region B encodes for five ORFs: B24, a fungal transcription factor (*STB4* ortholog); B25, a 5-oxoprolinase (*OXP1* ortholog); B26, a likely sugar-phosphate permease (D-galactonate transporter) or high-affinity nicotinic acid plasma membrane permease (*TNA1* ortholog); B27, a flocculin (containing a Flo11 domain and *FLO11* ortholog); and B28, a fungal transcription factor without an ortholog found (Novo, et al. 2009; Devia, et al. 2020). Interestingly, the putative functions of the ORFs identified within Region B do not necessarily imply a specific or exclusive role during fermentative processes. Accordingly, Region B presence is distributed along 14 different clades and not limited to Wine-European isolates (Legras, et al. 2018; O’Donnell, et al. 2023). Previous efforts to determine Region B structure and genome localization using draft genome assemblies identified a variable structure, in which Region B ORFs follow a circular order that suggests the presence of an extrachromosomal circular DNA (eccDNA), potentially facilitating its insertion into various chromosomal locations (Galeote, et al. 2011). Importantly, the genome sequencing of 1011 yeast isolates revealed a single strain carrying a longer Region B version within the genome (∼117 Kb), which suggests that Region B underwent a reduction in its genetic content (Peter, et al. 2018). Although most yeast strains carrying Region B contain the small version (∼17 Kb), the genetic structure and phenotypic contribution of this region across yeast strains remain largely undetermined.

In this work, we describe the horizontally transferred Region B at the genomic, transcriptional, and phenotypic levels. We used the ScRAP collection to characterize Region B at the genomic level, identifying a preference for chromosome (Chr) XI and a positioning in the core of chromosomes rather than subtelomeric regions. Then, we evaluated the transcriptional activity of each gene within Region B using five different genetic backgrounds not containing Region B. The results showed that ORFs within Region B are regulated by transcription factors encoded within this region, and the genetic background of the host strain. Finally, by performing Region B deletion in a wine yeast strain, we determined that Region B is involved in fermentation phenotypes such as fermentation rate and amino acids consumption. Altogether, our results demonstrate the molecular interaction between Region B and the host genome, contributing to yeast adaptation under fermentative conditions.

## 2. Results

### 2.1 Region B genomic structure suggests the presence of an extrachromosomal intermediate

To comprehensively analyze the structure of Region B through the *S. cerevisiae* genome evolution, multiple BLAST searches were performed for each ORF within Region B utilizing 142 yeast genomes from the ScRAP collection. Region B was identified 69 times across 39 yeast strains, spanning 15 of 29 phylogenetic clades (Supplementary Table S1; Fig. 1A), with a marked preference for Chr XI, and a location at the core chromosome rather than subtelomeric regions (defined according to the ScRAP collection information) (Fig. 1B, C, D). Then, we analyzed the Region B structure, observing ten distinct Region B variants, each displaying a different relative order of the ORFs, suggesting a circular continuity of the region (Fig. 2A, B). Given the unusual chromosomal positioning, variation in ORFs order, and structural diversity, these results suggest the possibility of an extrachromosomal circular DNA (eccDNA) intermediate, which facilitates the movement and integration of Region B into the genome (Fig. 2C). Among these variants, those containing all five ORFs were the most frequent, with six potential linearization points proposed for the hypothetical circular DNA (Fig. 2B). The remaining four less frequent variants may have originated from duplication events followed by ORF loss (Fig. 2A). Although Region B can reside in different chromosomes, some are exclusively —for instance, variant 10 on Chr X and variant 5 on Chr XV— localized in subtelomeric and core regions, respectively (Fig. 2A, C). In conclusion, Region B is widespread through clades, strains, and chromosomes in *S. cerevisiae*, with an unusual preference for the core of the chromosomes rather than subtelomeric regions. This region can be found in ten different variants, which strongly suggests the presence of an eccDNA as a possible intermediary for its integration and movement through the yeast genome.

**Figure 1.**
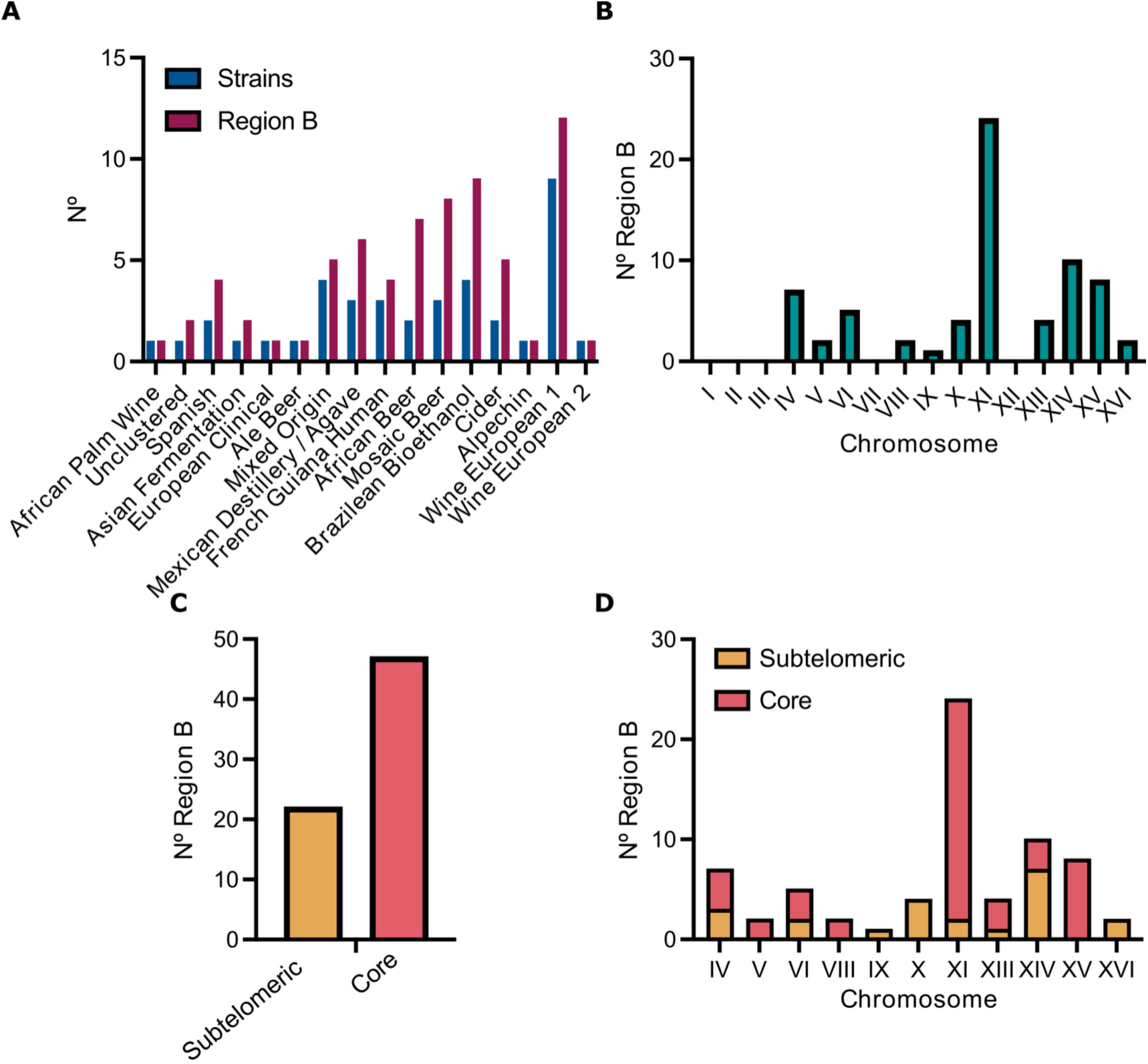
Region B is distributed in different *S. cerevisiae* clades with a preference for positioning in the central section (core) of the chromosomes. Region B was found across 15 of 29 clades of *S. cerevisiae*, with 69 regions B in 39 of 142 yeast strains of the ScRAP collection. (A) Number of yeast strains (blue) ordered by clades that harbor Region B and the number of Region B found in those strains (maroon). (B) Number of Region B on each yeast chromosome. (C) Position of Region B within chromosomes. (D) Position of Region B within each yeast chromosome.

**Figure 2.**
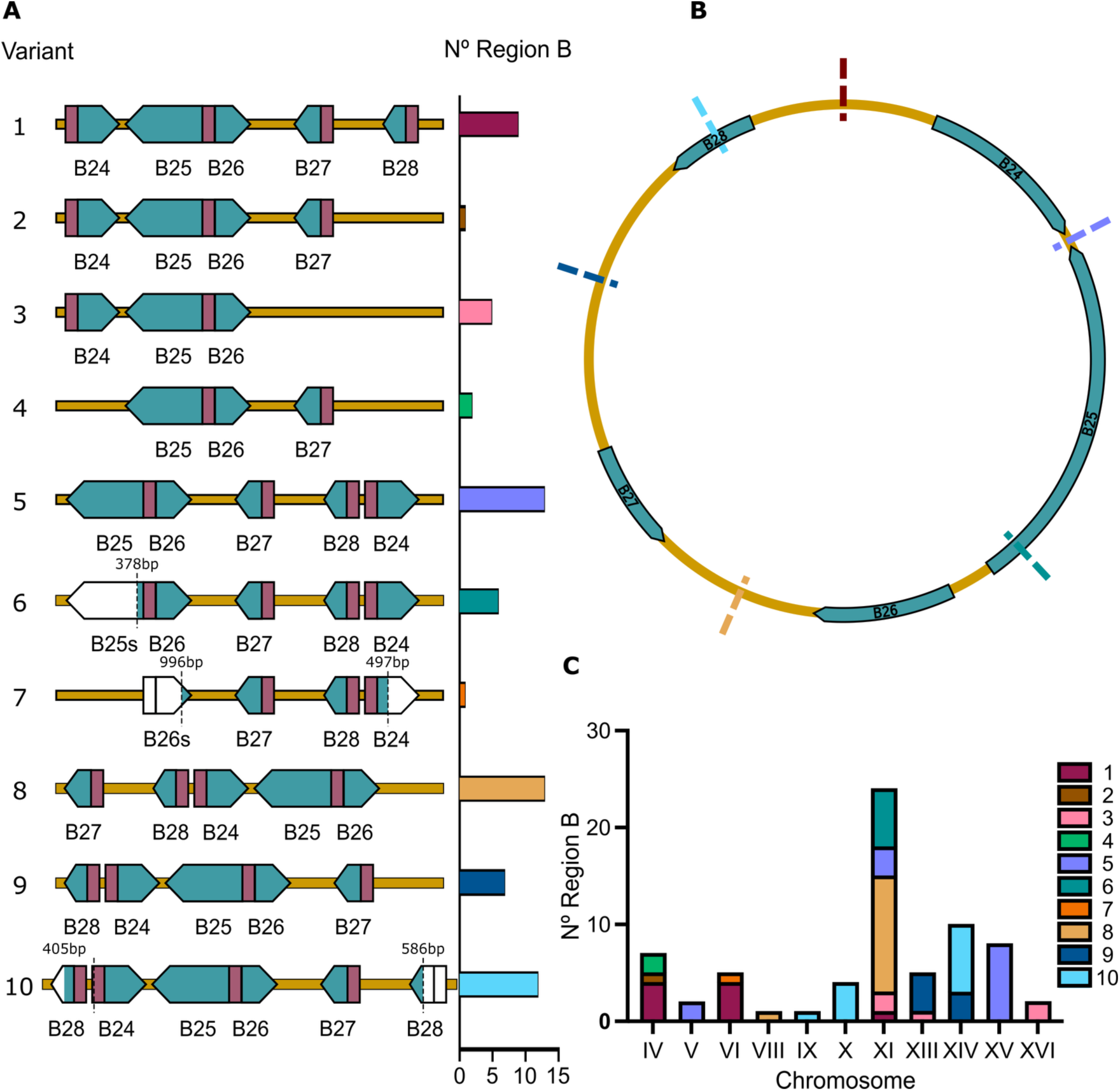
Region B structure in *S. cerevisiae* suggests the presence of an extrachromosomal intermediate. (A) 10 different Region B variants were identified in 39 strains of the ScRAP collection. Turquoise arrows represent the ORFs, maroon rectangles represent putative promoter regions, and white sections flanked by dashed lines indicate a truncated version of the ORF. Each variant is colored and quantified in the right panel. (B) A hypothetical circular intermediate of Region B showed 6 potential breakpoints for integration into the genome based on the most frequent variants. (C) The number of Region B was plotted according to the chromosome location of Region B variants.

### 2.2 Region B transcriptional activity is dependent on the host genetic background and its self-regulation

In yeast, HGT is often linked to domestication and fermentative traits, although the presence of Region B in yeast strains from diverse clades suggests broader functions. To explore this, 142 yeast strains from the ScRAP collection were phenotyped under seven growth conditions, comparing strains carrying (or not) Region B into the genome (Supplementary Fig. S1). Surprisingly, yeast strains carrying Region B exhibited higher growth in the Synthetic Complete (SC) medium with 1 mM hydrogen peroxide (H_2_O_2_) (Supplementary Fig. S1), suggesting a role in oxidative stress tolerance. As a control, we performed the same analysis, filtering the phenotypic data for comparison of yeast strains with the presence/absence of Region C in the genome (Supplementary Fig. S2). Interestingly, the results showed a growth decrease in SC galactose and growth at 40 °C for strains carrying Region C (Supplementary Fig. S2), suggesting a possible trade-off effect for this horizontally acquired region commonly associated with fermentation. Altogether, the ScRAP collection phenotyping suggests that Region B is implicated in yeast adaptation to oxidative stress.

Then, considering the phylogenetic distribution of Region B among different yeast clades, we hypothesized that the genetic background may influence the transcriptional activity of genes within this region. To address this, we selected a single yeast strain from the ScRAP collection for further analysis. This yeast strain was selected by extracting the pan-transcriptome information for the yeast strains of the ScRAP collection, identifying the AHG wine yeast strain as the one exhibiting the highest gene expression levels for genes within Region B (Fig. 3A). Thus, for each ORF within Region B, the promoter region from the AHG strain was used to control the expression of the *mCherry* reporter gene (Fig. 3B). This collection of five plasmids for transcriptional activity assays was transformed in five *S. cerevisiae* strains representing major genetic lineages: YPS128, North America (NA); Y12, Sake (SA), DBVPG6044, West African (WA); DBVPG6765, Wine European (WE); and the laboratory strains BY4741 (BY) as a control. Importantly, yeast strains selected from five different genetic background do not carry Region B encoded into their genomes (Fig. 4A). The transcriptional activity experiments showed almost no differences in *mCherry* expression for SA and BY genetic backgrounds compared with their controls, whereas NA, WA, and WE displayed higher *mCherry* expression with a variable timing (Fig. 4A). For instance, NA showed early expression (before 50 hours) for promoters from ORFs B25 (*pB25*), B26 (*pB26*), and B27 (*pB27*). In contrast, WA showed a delayed activation after 60 hours, and WE demonstrated early expression with increased activity around 70 hours (Fig. 4A). Altogether, the results indicate that Region B transcription is modulated by the host genetic background.

**Figure 3.**
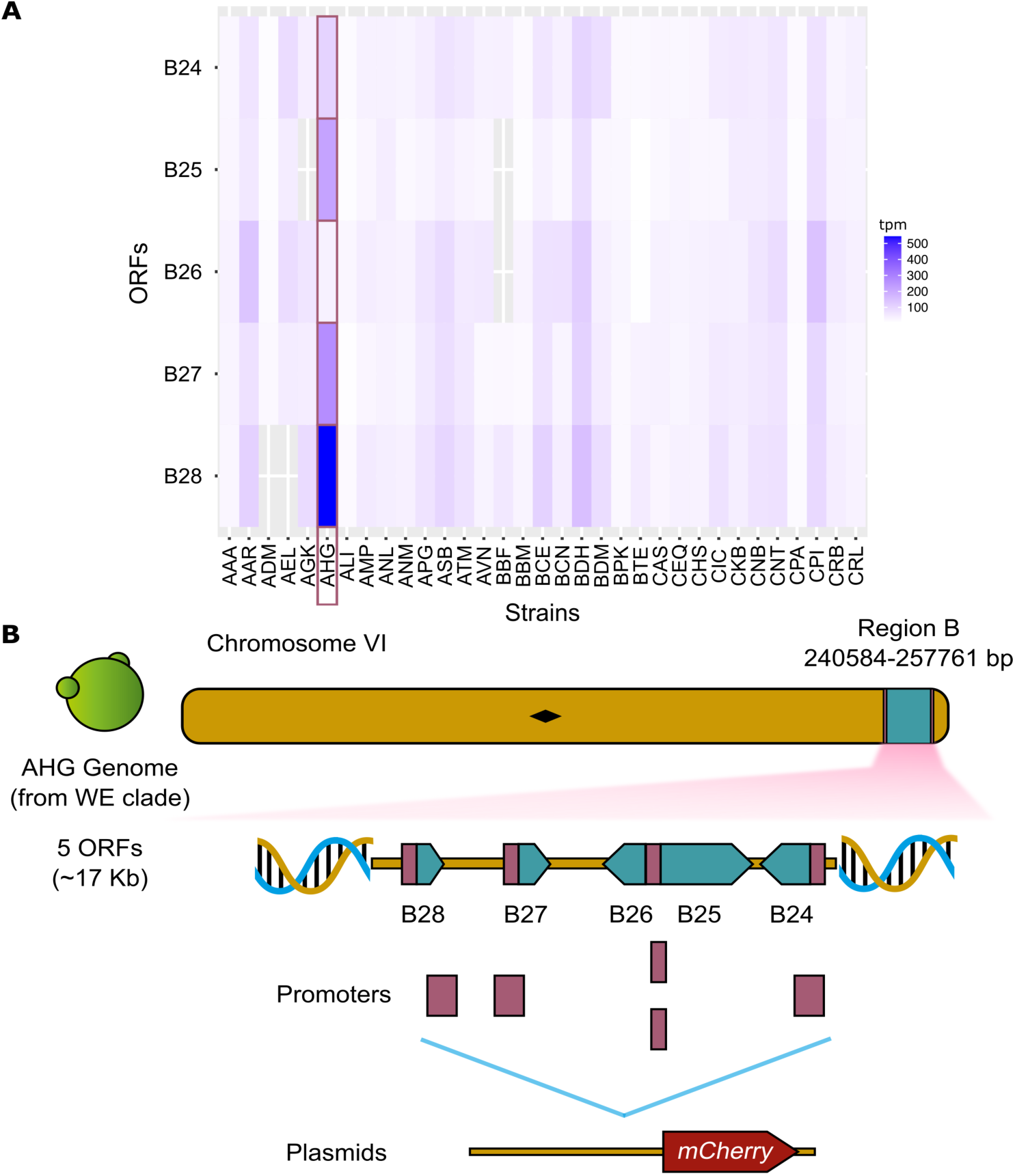
Selection of a single yeast strain from the ScRAP collection based on the pan-transcriptome information. (A) Normalized gene expression (Transcripts per million (tpm)) for each ORF within Region B in yeast strains from the ScRAP collection. The AHG wine yeast strain is highlighted in red. (B) Genetic structure of Region B in the AHG wine yeast strain. Promoter regions from each gene within Region B were used to evaluate the transcriptional activity, generating five different plasmids where each promoter region controls the expression of the *mCherry* reporter gene.

**Figure 4.**
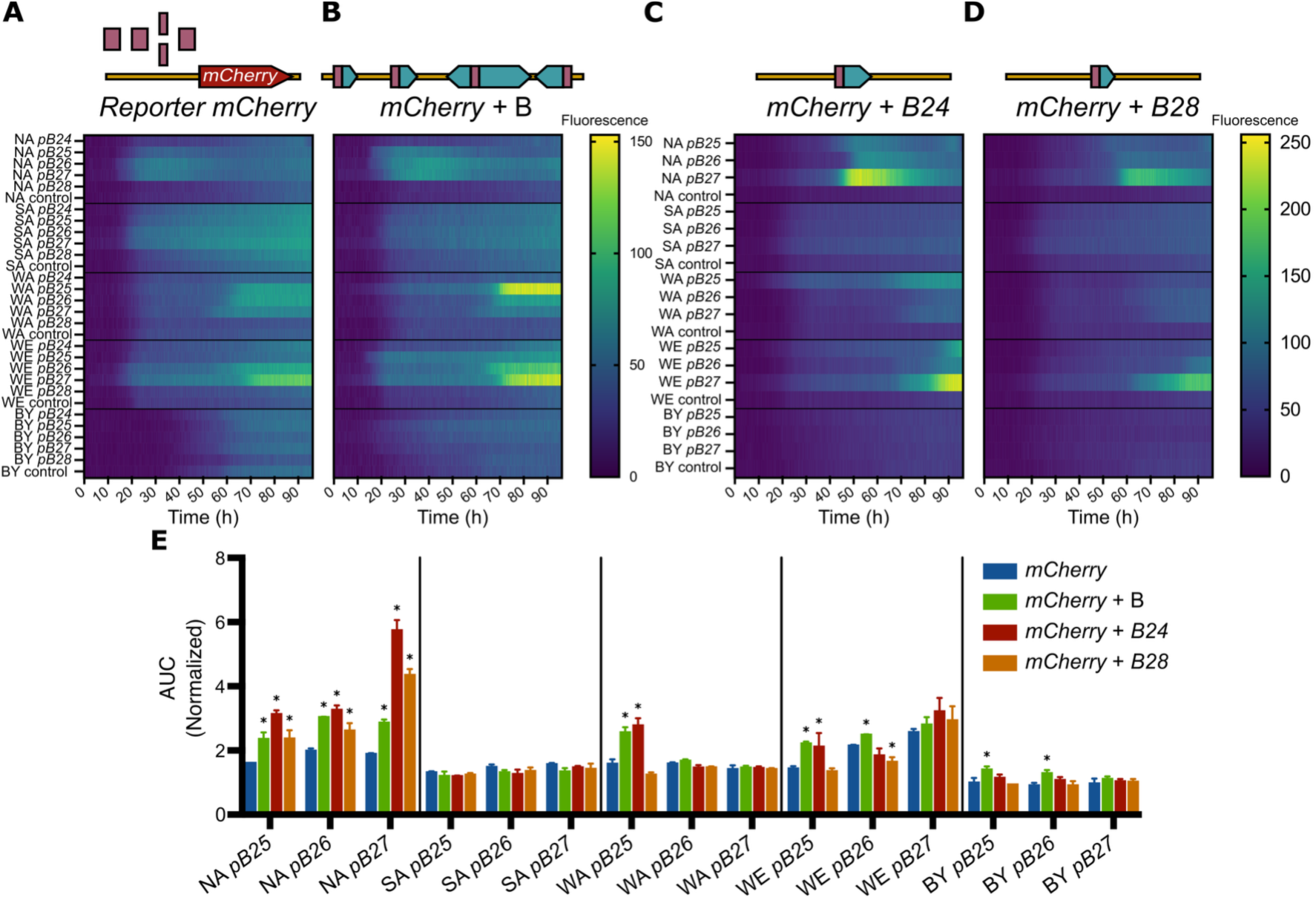
Transcriptional activity of Region B is affected by the host genetic background and its self-regulation. (A) Time course of transcriptional activity for each promoter from Region B (*pB24*, *pB25*, *pB26*, *pB27*, and *pB28*; represented as red boxes) in five different genetic backgrounds (NA, SA, WA, WE, and BY). *mCherry* expression was measured as fluorescence of the yeast cultures and represented as a heatmap. (B) Yeast strains shown in panel A co-transformed with a second plasmid encoding the entire region B. (C and D) Yeast strains shown in panel A co-transformed with a plasmid encoding the transcription factors B24 and B28, respectively. Heatmaps show the mean fluorescence every 10 minutes in 3 biological replicates. (E) Comparison of the transcriptional activity for different promoters and genetic backgrounds. Area Under the fluorescence Curve (AUC) was normalized by the fluorescence of the corresponding yeast strain carrying an empty vector, enabling comparison among experiments. Asterisks (*) indicate a statistically significant difference compared to the strain transformed only with the *mCherry* reporter (blue bar) plasmid (One-way ANOVA using Dunnett’s multi-comparison test, *p* < 0.05).

Region B encodes for two transcription factors (TFs), ORFs B24 and B28, that could be directly or indirectly regulating the expression of other ORFs within Region B. To uncover this, we co-transformed each strain with a second plasmid containing the entire Region B to evaluate a possible Region B self-regulation (Fig. 4B). These co-transformants showed similar temporal expression patterns but with higher expression levels in NA, WA, and WE genetic backgrounds (Fig. 4B, E). To elucidate the individual roles of each TF on other ORFs within Region B, yeast strains were co-transformed with plasmids encoding ORFs B24 or B28 independently (Fig. 4C, D). The results showed a higher level of expression but with a different temporal pattern (Fig. 4C, D, E). Remarkably, pB25 was transcriptionally active in WA and WE only when it was co-transformed with plasmids encoding either Region B or ORF-B24, suggesting a regulatory role for this TF. Overall, these results suggest that Region B regulates its transcription via TFs encoded within this region, but the timing and expression levels depend on the host’s genetic background.

### 2.3 Different genetic variants underlie Region B transcriptional regulation

To further investigate how the host genome regulates Region B expression, eQTLs were mapped, considering the temporal expression dynamics of *mCherry* transcriptional activity (Fig 4A). Thus, expression was classified as early (first two days) or late (last two days), in which NA (high early response) and WA (low early response) genetic backgrounds were selected to study early expression (Fig 4A). A previously described WAxNA hybrid strain and its derived collection of segregant strains were independently transformed with different reporter plasmids, including the promoter region from ORF-B25 (*pB25*), ORF-B26 (*pB25*), or ORF-B27 (*pB25*) controlling the *mCherry* reporter expression (Supplementary Fig. S3). After segregants phenotyping for transcriptional activity (*mCherry* expression; Supplementary Fig. S3), no significant eQTLs were found for *pB25* and *pB26* transcriptional activity, but two eQTLs were mapped for *pB27* expression (Chr V, position 86 Kb, LOD = 4.32, p = 0.001; Chr VII, position 142 Kb, LOD = 2.94, p = 0.036) (Fig. 5A). Then, to assess late expression, WE (high expression at later time points) and NA (low expression at later time points) were used (Fig. 4A). Similarly, the WExNA hybrid and its derived segregant strains were transformed with the *pB27* reporter plasmid and phenotyped (Supplementary Fig. S3). Interestingly, a different and more significant eQTL was mapped for this cross (Chr XII, position 420 Kb, LOD = 5.57, p = 0) (Fig. 5B).

**Figure 5.**
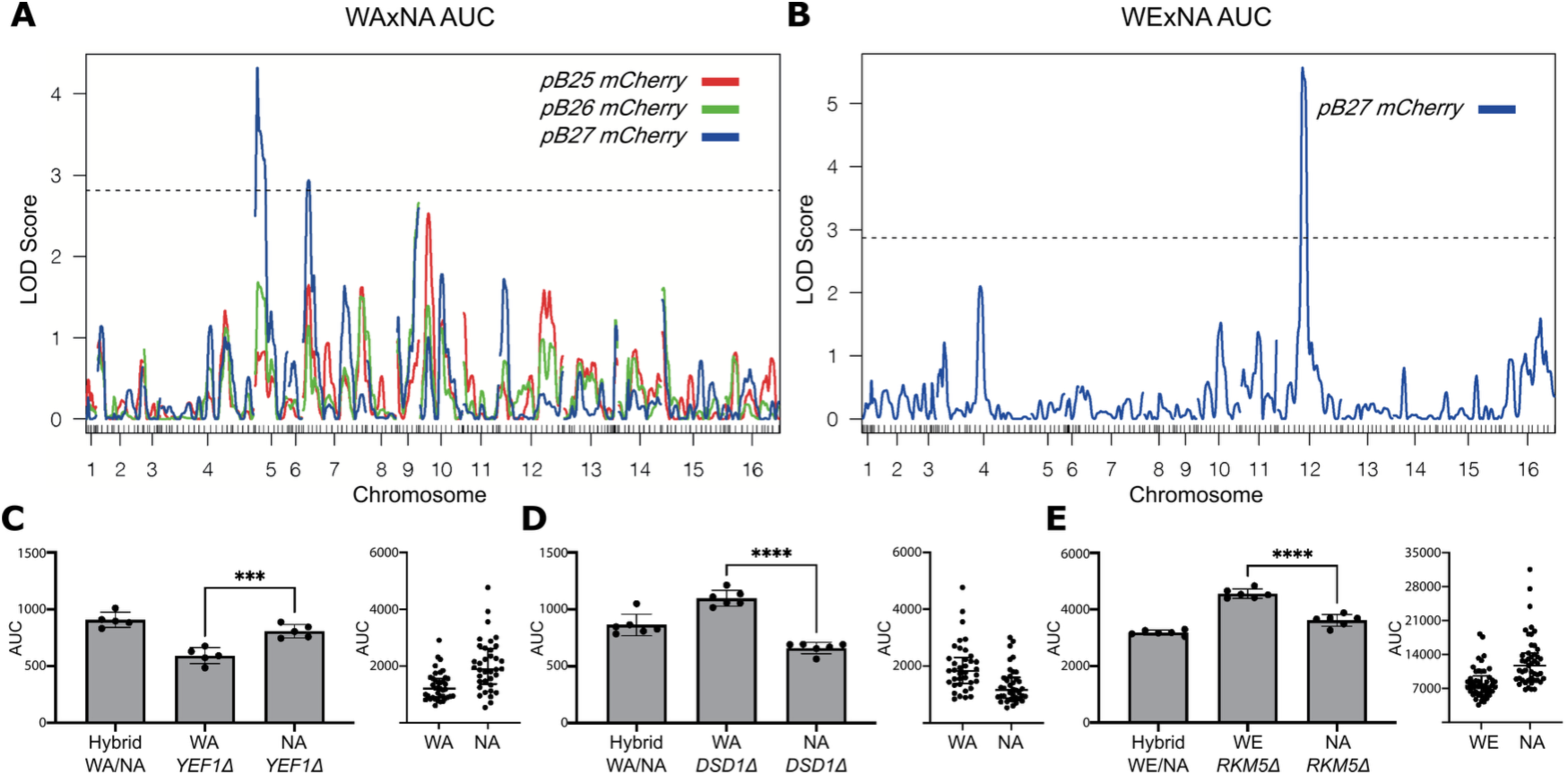
Different genetic variants affect the transcriptional activity of genes within Region B. (A and B) eQTLs mapped in the WAxNA cross (A) and WExNA cross (B) for transcriptional activity of the ORFs B25, B26, and B27. The promoter region of each ORF (pB25, pB26, and pB26) was fused to the *mCherry* fluorescent reporter. The dashed line represents the LOD significant threshold. (C, D, and E) Area Under the fluorescence Curve (AUC) for reciprocal hemizygous strains for *YEF1*, *DSD1,* and *RKM5* genes, respectively. The phenotype of the segregant strains and their respective genotypes at eQTL peak in each cross (WAxNA or WExNA) is shown in the right panel. Asterisks represent a statistically significant difference between hemizygous strains (Welch’s t-test, **** *p* < 0.0001, *** *p* < 0.001).

To validate the mapped eQTLs, candidate genes were selected within ±30 Kb of each QTL peak (Supplementary Table S2). Candidate genes were selected considering their described biological function, SNP number, and presence of non-synonymous SNPs. A total of 13 candidates were tested via reciprocal hemizygosity, resulting in 26 hemizygotes transformed with the *pB27* reporter plasmid (Supplementary Table S3). The results showed that *YEF1* (ATP-NADH kinase) and *DSD1* (D-serine dehydratase) genes accounted for 4.71% and 10.71% of the phenotypic variation observed in the WAxNA cross, respectively (Fig. 5C, D). Interestingly, both QTLs presented an antagonistic effect on *pB27* transcriptional activity (Fig. 5C, D), which suggests a complex transcriptional regulation. For eQTL mapped for the late transcriptional response, *RKM5* (a lysine methyltransferase) gene explained 9.15% of the phenotypic variation observed in the WExNA cross (Fig. 5E). Interestingly, the 3 validated genes indirectly regulate ORF-B27 promoter activity, since they do not encode for TFs (Supplementary Table S2). Despite this, our results suggest that horizontally acquired genes within Region B are transcribed depending on specific polymorphisms harbored by the host strains.

### 2.4 Region B contributes to yeast adaptation under fermentation conditions

To better understand the role of Region B in yeast adaptation, we performed the deletion of the entire region in the AHG strain (AHG *BΔ* strain). In addition, we generated a AHG strain with a deletion in the DNA segment encompassing the three central genes of Region B (ORF-B25, ORF-B26, and ORF-27), which are not encoding for transcription factors (AHG *EgΔ* strain). Then, we performed laboratory scale fermentations in SM60 (60 mg N/L; nitrogen limitation) and SM300 (300 mg N/L; nitrogen sufficiency) using the AHG wild-type strain (WT) and its derived mutants (*BΔ* and *EgΔ*). Fermentation progress was monitored as CO_2_ loss, extracting the maximal fermentation rate (Vmax) from the fermentation kinetics (Fig. 6A, B). Interestingly, in both fermentation conditions (SM60 and SM300), the AHG WT showed a higher Vmax compared to the AHG *BΔ* and *EgΔ* strains (Fig. 6A, B). Thus, results indicate that Region B contributes to maintaining the fermentation rate in the AHG wine yeast strain. Finally, we characterized the fermentation profile of each strain, quantifying by HPLC the production of different metabolites and the consumption of different amino acids (Supplementary Fig. S4, S5). In general, most phenotypic differences between yeast strains were observed in SM300, where nitrogen sufficiency enables fermentation progression and completion. Thus, a differential consumption of different amino acids was observed in the AHG BΔ and *EgΔ* strains respect to the AHG WT strain (Supplementary Fig. S6; Fig. 6C). Altogether, the results support the idea that Region B is implicated in amino acids consumption during the fermentation process.

**Figure 6.**
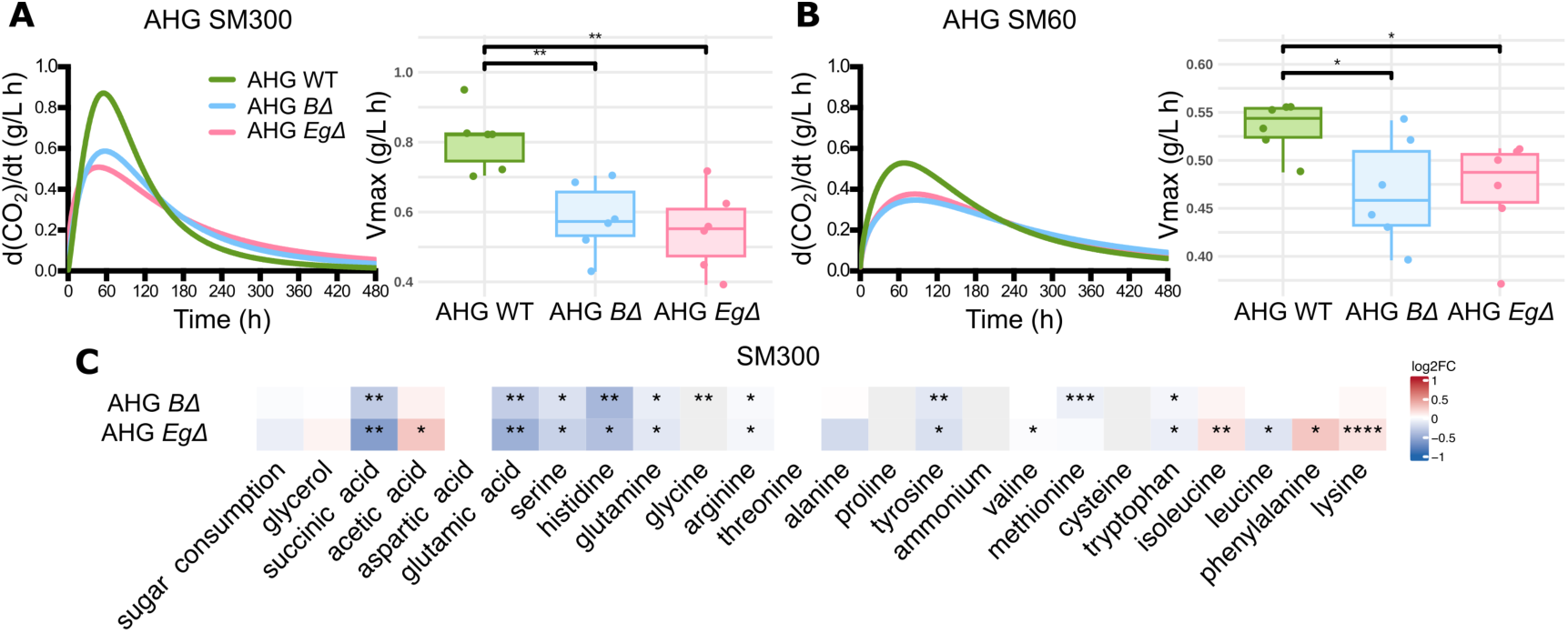
Region B is implicated in yeast adaptation to fermentative conditions. (A and B) Fermentation kinetics measured as CO_2_ loss curves (left) for the AHG strain and its derived strains in SM300 (A) and SM60 (B). The fermentation rate (Vmax) was extracted from the fermentation kinetics (right panel peak). Asterisks indicate statistically significant differences from the wild type (t-test, * p < 0.05, ** p < 0.01). (C) Heatmap highlighting the phenotypic differences in metabolites production and amino acids consumption for the AHG *BΔ* and AHG *EgΔ* strains respect to the wild-type strain (log_2_ fold change = 0) in SM300. In all panels, asterisks denote statistically significant differences respect to the WT phenotype determined by t-test or Wilcoxon test. Multiple testing correction was applied using the False Discovery Rate (FDR) method (* FDR < 0.05, ** FDR < 0.01, *** FDR < 0.001, **** FDR < 0.0001).

## 3. Discussion

Although HGT has traditionally been considered rare in eukaryotes, the increasing number of high-quality genome assemblies — facilitated by long-read technologies — has revealed that HGT continues to shape the eukaryotic web of life. In this study, we provide a comprehensive characterization of a horizontally acquired region (Region B) in *S. cerevisiae*, including its genomic distribution, transcriptional activity, and phenotypic contribution to yeast adaptation.

The genomic analysis of Region B posed technical challenges due to its structural variability across *S. cerevisiae* strains. To address this, a manually curated gene-by-gene BLAST search identified 11 additional genomes carrying Region B, increasing the number of genomes where Region B was previously detected in the ScRAP collection (O’Donnell, et al. 2023). Interestingly, Region B was predominantly located in chromosomal cores (47 instances) rather than subtelomeric regions (22 instances), contrasting with the typical subtelomeric localization of other horizontally acquired regions (Novo, et al. 2009). This unusual pattern may reflect a multi-step HGT event, possibly involving initial acquisition by *Z. parabailii* followed by transfer to *S. cerevisiae*, facilitated by Y’ elements and homologous recombination (O’Donnell, et al. 2023). Once inserted, Region B could be relocated within the yeast genome through mechanisms linked to Y’ mobility, such as unequal sister chromatid exchange or ectopic recombination (Louis, et al. 1994). In addition, retrotransposon-mediated processes — such as Ty1 LTR-based integration and the formation of virus-like particles (Louis and Haber 1992; Yamada, et al. 1998; Maxwell, et al. 2004) — could contribute to Region B patchy distribution. Notably, parallels exist in other eukaryotes, including Maverick elements in nematodes and recently described Starship elements in fungi, which facilitate gene cargo transfer and genome remodeling through virus-like strategies (Urquhart, et al. 2023; Widen, et al. 2023; Urquhart, et al. 2024). These observations underscore the evolutionary potential of modular mobile elements such as Region B and suggest the existence of similar mechanisms in yeast.

In *S. cerevisiae*, up to 23% of genomic content is implicated in the formation of extrachromosomal circular DNA (eccDNA), with LTR retrotransposons significantly overrepresented among these elements (Møller, et al. 2015). As proposed for Y’ elements, eccDNA can arise via intrachromatid homologous recombination and may serve as intermediates for reintegration, potentially explaining the patchy distribution of Region B across yeast strains. While the structural stability of core chromosomal regions may facilitate long-term retention and expression of Region B, in subtelomeric regions, its transcriptional activity could be repressed by sirtuins, particularly by Sir2 (Palladino, et al. 1993; Imai, et al. 2000; Wierman and Smith 2014). This silencing can limit the expression and functional integration of newly horizontally acquired genes. In contrast, insertions into core chromosomal regions may permit more robust transcription and a greater likelihood of functional incorporation, ultimately favoring retention through positive selection. Additionally, full-length variants of Region B (containing all five ORFs) were more frequently observed than truncated ones, which were often found co-occurring with at least one complete copy. This pattern supports the idea that partial duplications or rearrangements, possibly involving eccDNA intermediates, contributed to the diversity of Region B architectures. The higher frequency of Region B in chromosome XI remains an open question, warranting further investigation into chromosomal context and HGT integration dynamics.

The transcriptional activity of Region B was assessed by testing individual ORFs, their combinations, and the full-length region in different genetic backgrounds. These assays revealed striking differences in expression levels and timing depending on both the host strain and the presence of TFs encoded by Region B (Fig. 4). This suggests that Region B contains a self-regulatory module, although its activity remains modulated by the host genome context. Similar behavior has been described in horizontally acquired regions in other *Saccharomyces* species, where a horizontally transferred TF from *Zygosaccharomyces* regulates neighboring cargo genes, enabling coordinated expression (Tapia, et al. 2023). These findings support the idea that HGT loci can act as compact regulatory circuits, facilitating their functional integration into host networks. To further dissect the host’s influence, eQTL analysis was performed using a previously described segregant panel (Cubillos, et al. 2011). For the gene B27 (*FLO11* ortholog), three eQTLs were mapped and experimentally validated. Interestingly, none of the validated genes corresponded to canonical transcription factors, but rather to poorly characterized loci (Supplementary Table S2). This suggests that regulation of Region B expression may arise not from direct transcriptional control but from host-specific polymorphisms influencing the metabolic or epigenetic landscape. Such context-dependent regulation could allow horizontally transferred regions to remain transcriptionally silent under certain conditions and become active only when integration occurs into permissive genomic regions or the physiological environments are challenging. Altogether, these results highlight the dual nature of Region B regulation: TFs encoded within this region and the interaction with the host genome.

The effect of Region B on fermentative phenotypes was also determined, observing a direct implication in the fermentation rate (Vmax) under both SM300 and SM60 conditions (Fig. 6). Metabolites production and amino acids consumption during the fermentation process were also affected in the AHG *BΔ* and *EgΔ* strains compared to the WT strain (Fig. 6; Supplementary Fig. S6). Altogether, these results support the idea that Region B is functioning as a homeostatic regulatory module, regulating fermentation phenotypes under different nitrogen availability conditions.

Although more work is needed to fully understand the regulatory mechanisms of Region B, the predicted functions of its non-transcription factor genes (B25, B26, and B27) provide valuable insights. ORF-B25, which encodes a putative 5-oxoprolinase, has an ortholog linked to fungal development, stress responses, and host interaction in *Fusarium* species (Yang, et al. 2018), suggesting similar functions in *S. cerevisiae*. ORF-B26 likely encodes a nicotinic acid transporter, potentially increasing intracellular NAD^+^ levels and affecting glycolysis, amino acid metabolism, and lifespan-related traits such as telomeric silencing (Kato and Lin 2014; Orlandi, et al. 2020). These roles are consistent with the phenotypic and metabolic effects observed in this study. Finally, ORF-B27 encodes a Flo11-like adhesin, which may promote cell aggregation and changes in colony morphology, possibly mediating social behaviors or interactions with other *S. cerevisiae* strains or even *Z. parabailii* (Guo, et al. 2000; Chen and Fink 2006; Douglas Lois, et al. 2007; Váchová, et al. 2011; Brückner, et al. 2020). This suggests that Region B supports both physiological adaptation and cooperative strategies in shared environments.

In conclusion, our findings establish Region B as the most widely distributed horizontally acquired element in the *S. cerevisiae* pangenome. Despite its compact size, it exhibits transcriptional autonomy and regulatory plasticity, enabling it to interact with the host genome. These findings not only deepen our understanding of the yeast variable genome but also highlight the adaptive potential of horizontally acquired regions in optimizing fermentation performance.

## 4. Materials and Methods

### 4.1 Strains and culture conditions

The main yeast strains used in this work were: AHG (CBS1586); YPS128, North America (NA); Y12, Sake (SA), DBVPG6044, West African (WA); DBVPG6765, Wine European (WE); and the laboratory strains BY4741 (BY). The mutants, transformants, and segregants derived from the main strains are listed in the Supplementary Table S3. The strains were maintained in YPDA (2% glucose, 2% peptone, 1 yeast extract, and 2% agar) at 30 °C. Strains carrying plasmids with auxotrophic markers were maintained in Synthetic Complete (SC) medium (2% glucose, 0,67% YNB without amino acids, 0,2 dropout mix, and 2% agar) minus the corresponding amino acid or nucleotide (dropout mix).

### 4.2 Molecular cloning and strain construction

The genetic constructs generated in this work were assembled from PCR fragments using the *in vivo* yeast recombinational cloning method (Oldenburg, et al. 1997). For the reporter genetic constructs, promoter regions of each ORF within Region B were PCR-amplified from genomic DNA of the AHG strain, considering a promoter of 1000 bp or the intergenic region, while *mCherry* was used as reporter gene (Devia, et al. 2020). The overlapping PCR fragments were recombined into a pRS426 vector, where all the fragments were amplified using Phusion Flash High-Fidelity PCR master mix (Thermo Fisher Scientific, USA). The PCR products were co-transformed into the BY4741 yeast strain using the standard lithium acetate protocol (Gietz and Schiestl 2007). The assembled plasmids were extracted from yeast using the Zymoprep Yeast Plasmid Miniprep Kit (Zymo Research, USA). Plasmids were then transformed in *E. coli* (DH5α strain) and each fragment was confirmed by standard colony PCR. All the plasmids were confirmed by Sanger sequencing (Macrogen Inc., Republic of Korea). Plasmids and primers used and generated in this work are listed in the Supplementary Tables S4 and S5, respectively.

The same cloning procedure was used to generate plasmids encoding TFs B24 and B28 or the entire Region B. For the first case, terminators of the ORFs were considered as 300 bp or the intergenic region. For the plasmid encoding the entire region B, the region was divided into 3 fragments (5735 bp, 5834 bp, and 5788 bp), where each fragment was PCR amplified, enabling Region B assembly by yeast recombinational cloning (Oldenburg, et al. 1997). In both cases, we used the pRS315 vector to harbor the genetic constructs. Plasmids were transformed with standard lithium acetate protocol in different yeast strains and segregants strains derived from different crosses previously described (Cubillos, et al. 2011).

### 4.3 Individual gene deletion and Region B fragment deletion

Individual gene deletions were performed by one-step PCR amplification and direct homologous recombination with the following target genes: *LEU2, YEL025C, ECM10, YEF1, GTS1, IME4, GCN1, MDS3, TOS3, DSD1, ACE2, DPH6, RKM5,* and *GID11* (Baudin, et al. 1993). Briefly, the nourseothricin (*NatMx*) antibiotic resistance cassettes or *LEU2* gene were amplified by PCR using Phusion Flash high-fidelity PCR master mix (Thermo Scientific, Waltham, MA, USA) and 70 bp primers containing regions for direct homologous recombination with the target gene.

For deletion of the entire Region B (AHG *BΔ*) or a DNA segment within Region B encoding non-TFs (AHG *EgΔ*) ORFs (ORF-B25, ORF-B26, and ORF-B27) the CRISPR-Cas9 system was used in the AHG yeast strain, utilizing the pUDP004 plasmid encoding a single gRNA and the Cas9 protein (Gorter de Vries, et al. 2017). Briefly, a scarless repair was designed for each mutant, in which donor cassettes comprising the flanking 50 bp of each region of interest were synthesized as 100 bp oligos to repair the DNA. Different donor DNA used for CRISPR-Cas9 repair are listed in the Supplementary Table S6. The gRNAs were designed using the web tool CRISpy pop (https://crispy-pop.glbrc.org/) (Stoneman, et al. 2020). The gRNAs were inserted into a plasmid called pL58 derived from the pL59-Nat (Barre, et al. 2020), in which the nourseothricin (*NatMx*) antibiotic resistance was replaced by G418 resistance cassette (*KanMx*). The gRNAs were synthesized with the first six reverse complement bases of the gRNA for splicing with the hammerhead ribozyme and inserted into the *BsaI* site of the pL58 plasmid by Gibson assembly (Gibson, et al. 2009). The final plasmids for CRISPR-Cas9 deletion were transformed in the AHG strain using the lithium acetate protocol (Gietz and Schiestl 2007).

### 4.4 RNA-seq data analysis and *mCherry* expression

RNA-seq data from the *S. cerevisiae* pan-transcriptome (Caudal, et al. 2024) was utilized to analyze the expression levels of yeast strains carrying region B into the genome, but focused on the 142 telomere-to-telomere genomes of the ScRAP collection (O’Donnell, et al. 2023). Briefly, data available in the European Nucleotide Archive (https://www.ebi.ac.uk/ena/browser/home) was filtered using the 5 genes within Region B: B24 (X17-EC1118_1N26_0056g), B25 (X13-EC1118_1N26_0012g), B26 (X14-EC1118_1N26_0023g), B27 (X15-EC1118_1N26_0034g), and B28 (X16-EC1118_1N26_0045g). Then, data was filtered using the names of the 39 yeast strains from the ScRAP collection carrying the Region B. This allowed the extraction of the transcripts per million (tpm) for each strain, which was finally graphed as a heatmap for each ORF of Region B.

For *mCherry* reporter gene expression, we used our previously described protocol for simultaneous determination of optical density and fluorescence of the yeast cultures (Devia, et al. 2020; Figueroa, et al. 2025; Ruiz, et al. 2025). Briefly, each yeast strain carrying a reporter plasmid was grown overnight in a 96-well plate containing 200 μL of YNB media. Then, 15 μL of the overnight cultures were transferred to a new plate with optical bottom (ThermoFisher Scientific, USA) containing 285 μL of YNB media. Optical Density at 600 nm (O.D._600_) and *mCherry* fluorescence were measured every 10 minutes for 4 days (96 h) using a Synergy H1M plate reader (Agilent, USA) at 25 °C. Area Under the Fluorescence Curve (AUC) was quantified using GraphPad Prism version 10.1.1 (Dotmatics, USA), enabling the determination of the total fluorescence in the yeast culture (Ruiz, et al. 2025). Fluorescence experiments were performed in three biological replicates.

### 4.5 QTL mapping and validation

Linkage analysis between phenotype (fluorescence AUC) and genotype was performed with R/qtl software (Broman, et al. 2003), using a non-parametric model to determine the LOD scores. A 0.05 tail with 1000 permutations was used to determine the significance of a QTL (Cubillos, et al. 2011). The genotype of the segregant strains from the WAxNA and WExNA crosses was previously described (Cubillos, et al. 2011; Salinas, et al. 2012).

For QTLs validation, a reciprocal hemizygosity analysis was performed (Steinmetz, et al. 2002). Hemizygous yeast strains were obtained by deletion of the candidate gene with the *LEU2* gene and using the parental yeast strains (WA, NA, and WE) with the following genotype: *MAT* a, *ura3::KanMx*, *leu2::NatMx*. These strains were crossed with the parental yeast strain carrying the wild type allele of interest: *MAT* α, *ura3::KanMx*. Then, deletion of the candidate gene in the opposite mating type (*MAT α*) was also performed and this strain was crossed with the parental strain carrying the wild-type allele (*MAT* a). Thus two isogenic yeast strains that differed in a single allele were generated, whose phenotypes were compared.

### 4.6 ScRAP collection phenotyping

Quantitative phenotyping was performed using an end-point colony growth determination on solid Synthetic Complete (SC) media with a different stressors or carbon sources. The ScRAP collection strains were pre-grown on solid YPD using the replicating ROTOR HDA benchtop robot (Singer Instruments, England), starting from a glycerol stock of two 96-well microplates. Seven conditions were assessed in a solid matrix of 384 spots from the YPD plates, in which each strain was evaluated in quadruplicates on the corresponding matrix. The plates were incubated at 30 °C (except for the condition at 40 °C) for 2 days in the assays of Fluconazole 25 μg/mL, temperature 40 °C, and NaCl 1.5 M; and 3 days for Galactose 2%, Glycerol 2%, and SC (control) conditions. Then, plates were scanned with a resolution of 600 dpi and 16-bit grayscale. The quantification of colony size from the plate images was performed using SGAtools (Wagih, et al. 2013), and the values were normalized per plate based on row/column medians. Finally, the relative growth was quantified for each strain as the ratio of the medians in the condition of interest and YPD.

### 4.7 Fermentations conditions

In the fermentation experiments yeast strains were growth in SM60 (60 mg N/L) and SM300 (300 mg N/L) cultures media containing different amounts of Yeast Assimilable Nitrogen (YAN), which were prepared according to recipes previously described (Kessi-Pérez, et al. 2019; Molinet, et al. 2020). Then, laboratory scale fermentations in SM60 and SM300 were performed according to previously described protocols (Kessi-Pérez, et al. 2019; Molinet, et al. 2020). Briefly, yeast strains were pre-grown in SM60 and SM300 for 48 h at 28 °C without shaking. Then, 15 mL tubes containing 12 mL of SM60 or SM300 were inoculated with 10^6^ cells per mL and incubated at 25°C for 20 days without agitation. Fermentations were monitored weighing the fermentation tubes daily and determining weight loss over time (Kessi-Pérez, et al. 2019; Molinet, et al. 2020). The CO_2_ loss curves were obtained by fitting the weight loss data to a sigmoid non-linear regression using the GraphPad Prism software version 10.1.1 (Dotmatics, USA). From the fitted CO_2_ curves, the fist derivative was calculated to obtain the maximal fermentation rate (Vmax) of each strain using RStudio (2024.04.2+764). All the fermentation experiments were carried out in six biological replicates.

At the end of the fermentations, production of different metabolites was carried out by HPLC, including glycerol, acetic acid, succinic acid, and residual sugar (glucose and fructose). For this, 50 µL of filtrated must was injected in a Shimadzu Prominence HPLC equipment (Shimadzu, USA) using a Bio-Rad HPX –87H column (Nissen, et al. 1997). Similarly, amino acids were determined by HPLC with a previous derivatization process with diethyl ethoxymethylenemalonate (DEEMM) (Gómez-Alonso, et al. 2007). The amino acids consumption was determined as the difference between the initial and final concentrations before and after fermentation, respectively (Molinet, et al. 2019; Molinet, et al. 2020).

## 5. Acknowledgments

This research was funded by the ANID-Millennium Science Initiative Program-ICN17_022 to F.A.C. and F.S.; the ANID-FONDECYT grant number 1210955 to F.S.; the ANID-FONDECYT grant number 1220026 to F.A.C.; the ANID-PhD scholarships 21210525 to A.R.; ANID-Subdirección de Investigación Apicada grant number ID24I10027 to E.K.P.; ANID-FONDECYT grant number 1250815 to C.M. We thank Diego Ruiz for technical help during this project execution.

## 6. Author Contributions

Conceptualization, A.R. and F.S.; methodology, A.R., C.B., M.DC., X.S. H.C., and B.B.; analysis, A.R., C.B., M.DC., X.S. H.C., and B.B.; investigation, A.R., C.B., M.DC., X.S. H.C., and B.B.; data curation, A.R. and F.S.; writing original draft preparation, A.R. and F.S.; writing review and editing, F.S.; supervision, F.A.C., C.M., E.I.KP., G.L. and F.S.; funding acquisition, F.A.C., C.M., E.I.KP., G.L. and F.S. All authors have read and approved the final version of the manuscript.

