## Supplementary Figures for "Assessing the molecular and phenotypic contribution of a horizontally acquired region to yeast adaptation"

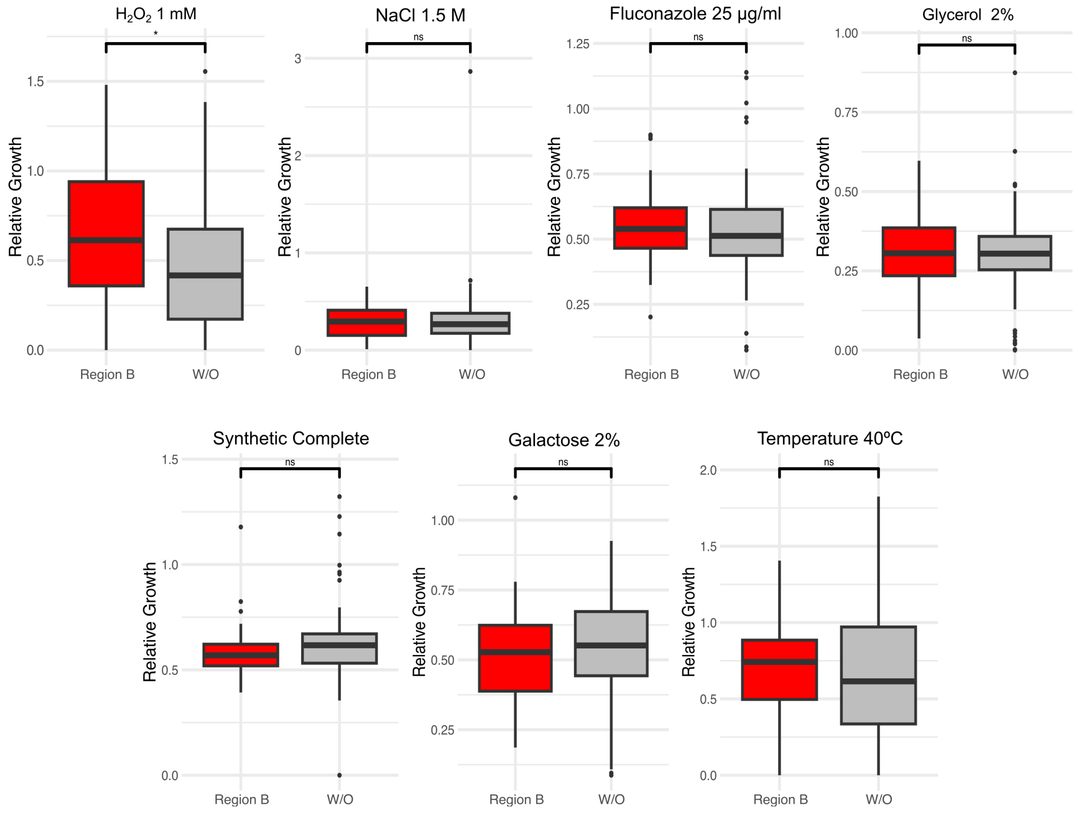


**Supplementary Figure S1**. Growth quantification in yeast strains from the ScRAP collection carrying (or not) the region B. Growth differences in yeast strains carrying Region B (n=39) and without (W/O) this region (n=102) were assayed in different culture conditions. No statistical differences were found between the two sets of yeast strains.

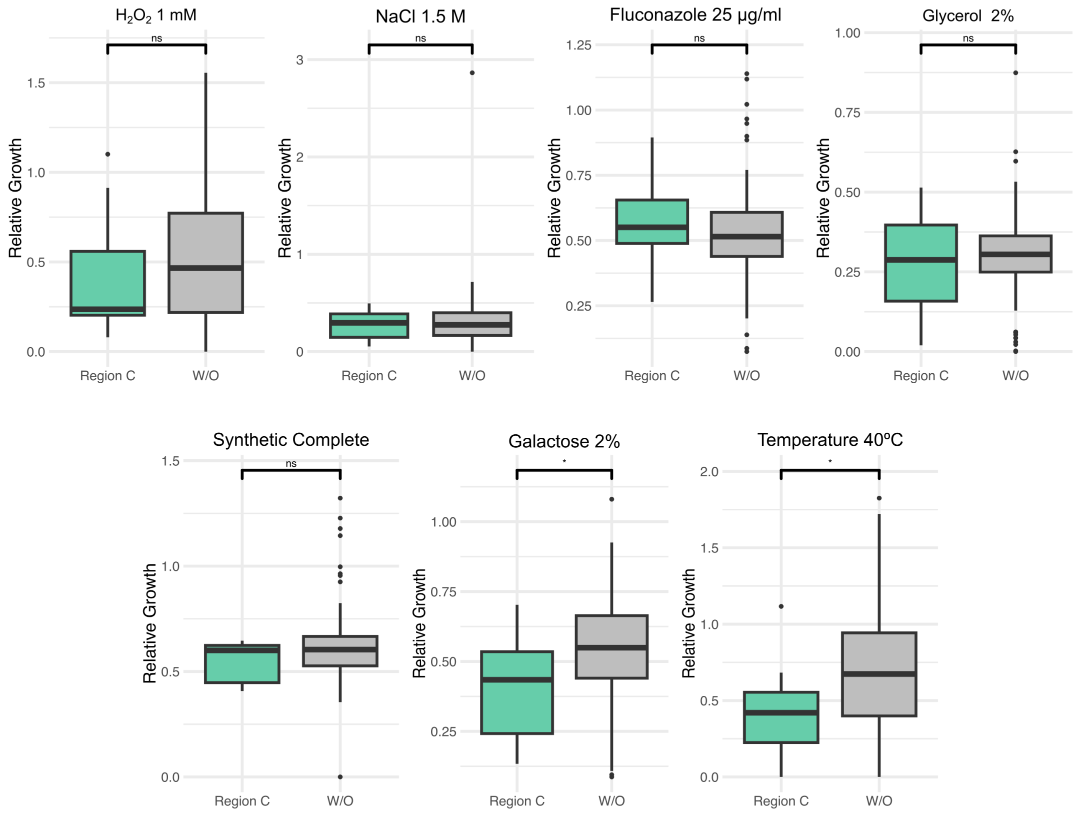


**Supplementary Figure S2**. Growth quantification for yeast strains from the ScRAP collection carrying (or not) Region C within the genome. Growth differences between yeast strains carrying Region C (n=9) compared to yeast strains without (W/O) the region (n=132) in the genome were assayed in different growth conditions. Asterisk (*) represents a statistically significant difference between the two groups of yeast strains (Mann-Whitney test, $p < 0.05$).

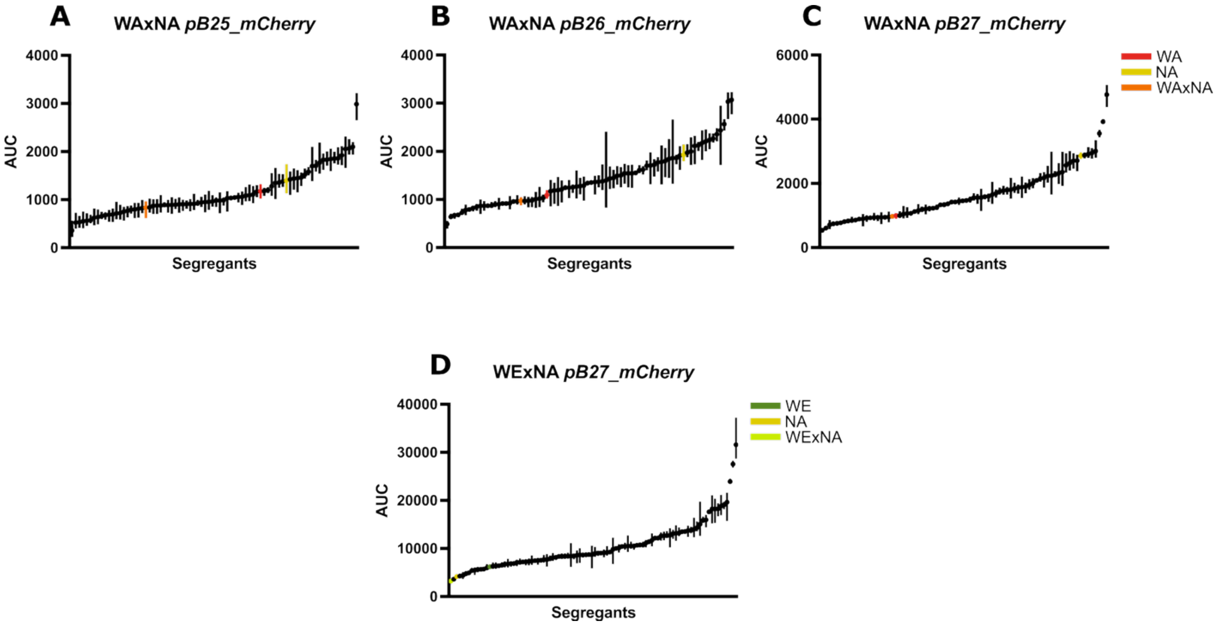


**Supplementary Figure S3**. Transcriptional activity for genes within Region B in segregant strains from the WAxNA and WExNA crosses. *mCherry* fluorescence, measured as Area Under the fluorescence Curve (AUC), was plotted in the y-axis for 75 segregants from the WAxNA cross, which were transformed with the reporter plasmids for: (A) ORF-B25 (*pB25*), (B) ORF-B26 (*pB26*), and (C) ORF-B27 (*pB27*) promoters. (D) *mCherry* fluorescence was measured as AUC for 93 segregants from the WExNA cross and transformed with the reporter plasmid for the ORF-B27 (*pB27*) promoter. The phenotypes of the parental strains and their hybrids are indicated with the color code.

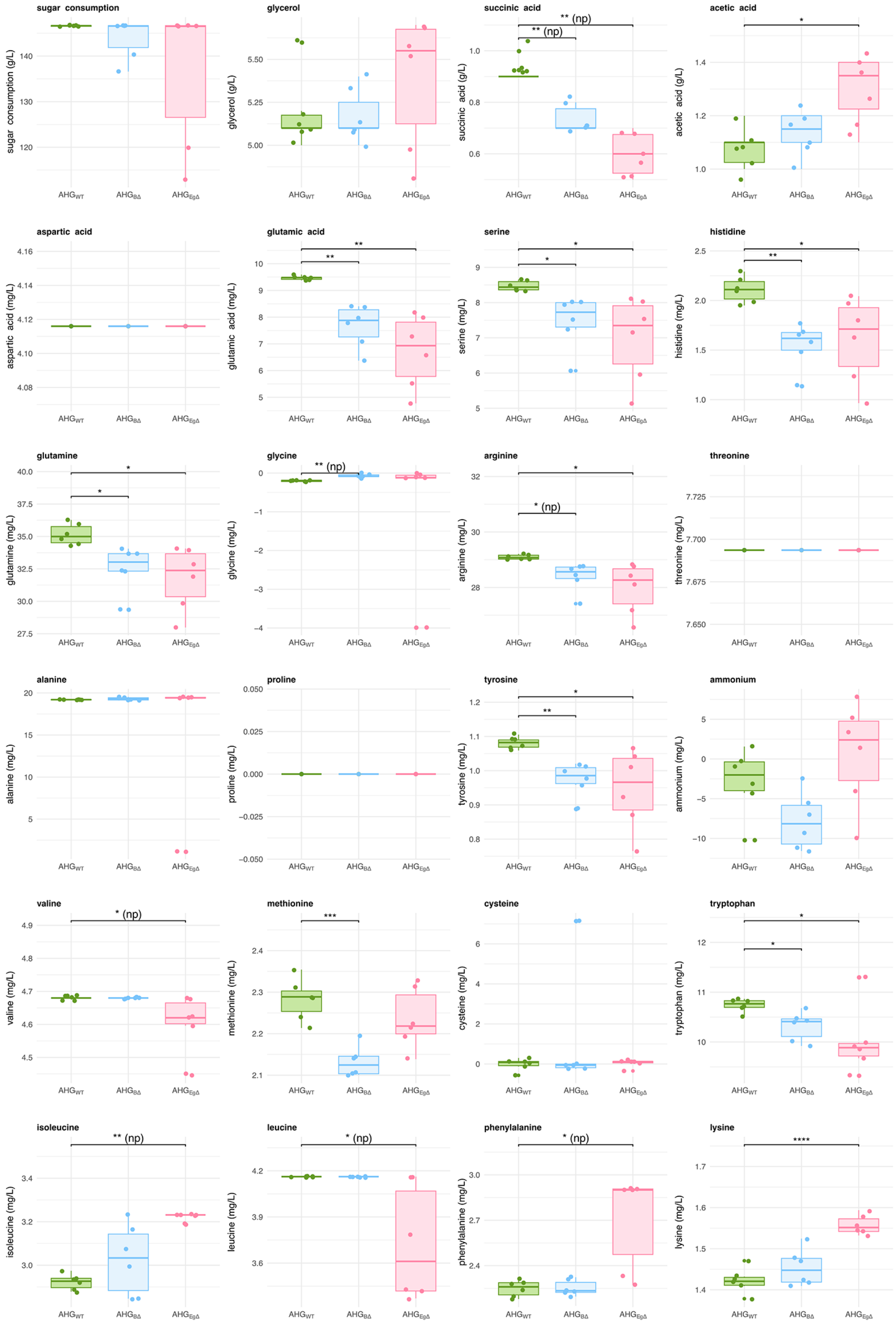
sugar consumption
sugar consumption (g/L)
glycerol
glycerol (g/L)
succinic acid
succinic acid (g/L)
** (np)
** (np)
acetic acid
acetic acid (g/L)
*
aspartic acid
aspartic acid (mg/L)
glutamic acid
glutamic acid (mg/L)
**
**
serine
serine (mg/L)
*
*
histidine
histidine (mg/L)
*
**
glutamine
glutamine (mg/L)
*
*
glycine
glycine (mg/L)
** (np)
arginine
arginine (mg/L)
*
* (np)
threonine
threonine (mg/L)
alanine
alanine (mg/L)
proline
proline (mg/L)
tyrosine
tyrosine (mg/L)
*
**
ammonium
ammonium (mg/L)
valine
valine (mg/L)
* (np)
methionine
methionine (mg/L)
***
cysteine
cysteine (mg/L)
tryptophan
tryptophan (mg/L)
*
*
isoleucine
isoleucine (mg/L)
** (np)
leucine
leucine (mg/L)
* (np)
phenylalanine
phenylalanine (mg/L)
* (np)
lysine
lysine (mg/L)
****
$AHG_{WT}$
$AHG_{BΔ}$
$AHG_{EgΔ}$

**Supplementary Figure S4.** Fermentation by-products and nitrogen source consumption in AHG wild type and its derived strains (AHG *BΔ* and AHG *EgΔ*) in SM300. HPLC quantification of the metabolites was performed after 20 days of fermentation. Asterisks denote statistically significant differences determined by t-test or Wilcoxon test, while np denotes non-parametric data. Multiple testing correction was applied using the False Discovery Rate (FDR) method (* FDR < 0.05, ** FDR < 0.01, *** FDR < 0.001, **** FDR < 0.0001).

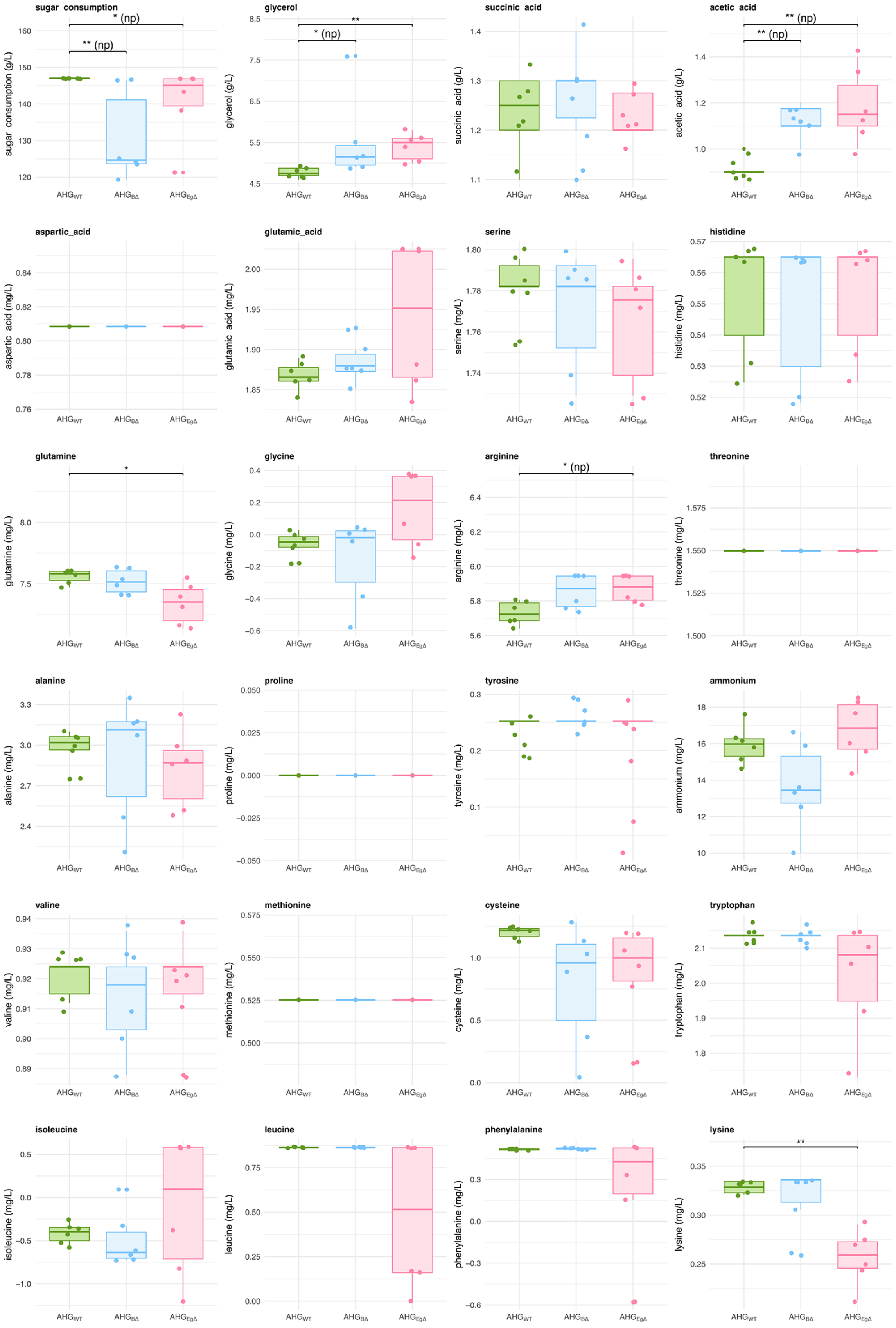
sugar consumption
sugar consumption (g/L)
** (np)
* (np)
glycerol
glycerol (g/L)
* (np)
**
succinic acid
succinic acid (g/L)
acetic acid
acetic acid (g/L)
** (np)
** (np)
aspartic_acid
aspartic acid (mg/L)
glutamic_acid
glutamic acid (mg/L)
serine
serine (mg/L)
histidine
histidine (mg/L)
glutamine
glutamine (mg/L)
*
glycine
glycine (mg/L)
arginine
arginine (mg/L)
* (np)
threonine
threonine (mg/L)
alanine
alanine (mg/L)
proline
proline (mg/L)
tyrosine
tyrosine (mg/L)
ammonium
ammonium (mg/L)
valine
valine (mg/L)
methionine
methionine (mg/L)
cysteine
cysteine (mg/L)
tryptophan
tryptophan (mg/L)
isoleucine
isoleucine (mg/L)
leucine
leucine (mg/L)
phenylalanine
phenylalanine (mg/L)
lysine
lysine (mg/L)
**
AHG_WT
AHG_BΔ
AHG_EgΔ

**Supplementary Figure S5.** Fermentation by-products and nitrogen source consumption in AHG wild type and its derided strains (AHG *BΔ* and AHG *EgΔ*) in SM60. HPLC measured the metabolites after 20 days of fermentation. Asterisks indicate statistically significant differences identified by t-test or Wilcoxon test, while np denotes non-parametric data. Multiple testing correction was applied using the False Discovery Rate (FDR) method (* FDR < 0.05, ** FDR < 0.01, *** FDR < 0.001, **** FDR < 0.0001).

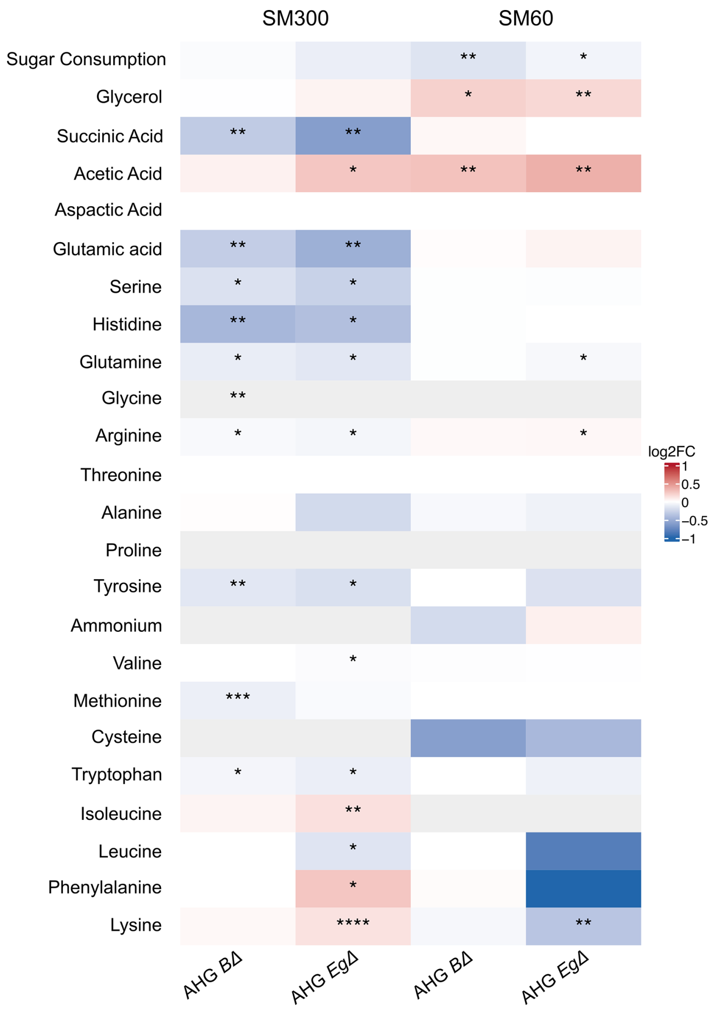


**Supplementary Figure S6.** Phenotypic differences for the AHG strain carrying different Region B deletions compared to the wild-type strain. Heatmap showing the $\log_2$ fold change (log2FC) for the fermentative by-products and nitrogen consumption (amino acids) in the AHG mutant strains (*BΔ* and *EgΔ*) respect to the wild-type phenotype in SM300 and SM60. Values represent changes relative to the wild type strain ($\log_2$ fold change = 0). Asterisks denote statistically significant differences determined by t-test or Wilcoxon test. Multiple testing correction was applied using the False Discovery Rate (FDR) method (* FDR < 0.05, ** FDR < 0.01, *** FDR < 0.001, **** FDR < 0.0001).
